# *In silico* study predicts a key role of RNA-binding domains 3 and 4 in nucleolin-miRNA interactions

**DOI:** 10.1101/2021.06.09.447752

**Authors:** Avdar San, Dario Palmieri, Anjana Saxena, Shaneen Singh

## Abstract

RNA binding proteins (RBPs) regulate many important cellular processes through their interactions with RNA molecules. RBPs are critical for post-transcriptional mechanisms keeping gene regulation in a fine equilibrium. Conversely, dysregulation of RBPs and RNA metabolism pathways is an established hallmark of tumorigenesis. Human nucleolin (NCL) is a multifunctional RBP that interacts with different types of RNA molecules, in part through its four RNA binding domains (RBDs). Particularly, NCL interacts directly with microRNAs (miRNAs) and is involved in their aberrant processing linked with many cancers, including breast cancer. Nonetheless, molecular details of the NCL-miRNA interaction remain obscure. In this study, we used an *in silico* approach to characterize how NCL targets miRNAs and whether this specificity is imposed by a definite RBD-interface. Here, we present structural models of NCL-RBDs and miRNAs, as well as predict scenarios of NCL- miRNA interactions generated using docking algorithms. Our study suggests a predominant role of NCL RBDs 3 and 4 (RBD3-4) in miRNA binding. We provide detailed analyses of specific motifs/residues at the NCL- substrate interface in both these RBDs and miRNAs. Finally, we propose that the evolutionary emergence of more than two RBDs in NCL in higher organisms coincides with its additional role/s in miRNA processing. Our study shows that RBD3-4 display sequence/structural determinants to specifically recognize miRNA precursor molecules. Moreover, the insights from this study can ultimately support the design of novel antineoplastic drugs aimed at regulating NCL-dependent biological pathways with a causal role in tumorigenesis.

**Importance/impact of the study:** Nucleolin is a multifunctional RNA binding protein that is often linked with many cancers. Similarly, microRNAs are often dysregulated in many cancers and linked to tumorigenesis. This study focuses on the interaction of nucleolin with microRNAs to identify previously unknown mechanistic details/specificity of these interactions. The insights from this study can ultimately support the design of novel drugs aimed at regulating NCL-dependent pathways implicated in tumorigenesis.

## Introduction

RNA binding proteins (RBPs) are critical in modulating RNA metabolism and linked with erroneous gene regulation in a wide range of disease conditions^1^. The human genome codes for more than 3500 RBPs^2^. Their emerging role is underscored by genome-wide studies indicating that hundreds of these RBPs are significantly dysregulated in a variety of cancer types^3^, where some are even identified as potential cancer drivers^4^. In addition, RBPs are implicated in numerous somatic and mendelian genetic diseases, impacting multiple organ systems in humans such as metabolic, neurodegenerative, musculoskeletal and connective tissue diseases^2^. Altered expression or function of RBPs translates into aberrant control of target RNAs, and hence gene expression, ultimately driving pathological phenotypes^5^. RBP-RNA interactions are driven by RNA-binding domains (RBDs) and are often dysregulated in human cancers^5^. Importantly, many RBPs bind the same sub-set of target RNAs, potentially exploiting a synergistic or competitive physiology^6^. However, the molecular mechanisms by which RBPs direct their specificity, including the selective use of their constituent RBDs to target specific RNA types, remain elusive. RBPs are known to interact with RNA molecules through two RNA-binding motifs (RNP1 and RNP2). RNPs within an RBD provide some underlying principles about how an RBP recognizes specific RNA species^7^. RNPs are evolutionarily conserved among many RBPs and correspond to a β_1_-α_1_-β**_2_-**β**_3_**-α_2_-β_4_ structural arrangement . The two beta strands found in the middle of this arrangement (indicated in bold) are known to interact with RNA either as the octameric RNP1 or as the hexameric RNP2 motif with the conserved sequence (R/K)-G-(F/Y)- (G/A)-(F/Y)-V-X-(F/Y) [10] or as the hexameric RNP2 motif with the conserved sequence of (L/I)-(F/Y)-(V/I)-X-(N/G)-L^9^. RNP-RNA interactions are predominantly hydrophobic, and aromatic residues are especially important in mediating the interaction through Van der Waals forces, π-π stacking interactions10 with nucleotide bases and π-sugar ring interactions^10^. Additionally, basic residues in these conserved motifs also form salt bridges with phosphate groups to enhance stability^10^. Previous studies have established the role of aromatic and basic residues in interactions of hnRNP A1^11–12^ and Lin28^13^ with RNA molecules.

Nucleolin (NCL), a multifunctional RBP is often overexpressed in many cancers and disease conditions^14^. NCL is involved in myriads of cellular processes that are ultimately tied to its RNA/DNA-binding functions to regulate gene expression that control cell survival, growth and or death. These roles include sensing stress^15^, ribosome biogenesis^16^, chromatin remodeling^17^, DNA replication, transcription, messenger RNA (mRNA) turnover^18^, induction^19^ & inhibition^20^ of translation, and microRNA (miRNA) biogenesis^21^. NCL protein is organized into distinct functional domains: (a) the highly acidic N-terminal domain with basic stretches that contains the nuclear localization signal, is heavily phosphorylated during the cell cycle by stage-specific kinases, and drives the histone chaperone activity of NCL^17^; (b) the glycine and arginine-rich (RGG/GAR) C-terminal domain, known to play a critical role in protein-protein interactions such as with ribosomal proteins^22^ and the tumor suppressor p53^23^ and is also implicated in non-specific interactions with RNA; and (c) the central region constitutes two-to-four distinct RNA-binding domains and is critical for its interaction with different species of RNAs^24^. Most eukaryotic species, including plants, contain only two RBDs in NCL protein, where the individual RBD domains are better conserved among the orthologs than within the protein. Interestingly, NCL from *Dictyostelium Discoideum* uniquely possesses an odd number of RBDs (three RBDs) suggesting a unique RNA binding profile in this organism (Singh Lab, unpublished). NCL has evolved in vertebrates, including humans, to an increased (four) number of RBDs where RBDs 3 and 4 are unique to these organisms^24^. It is also well-established that RBDs 1 and 2 are sufficient for certain NCL-RNA interactions, specifically binding to mRNA^25, 26^ and rRNA molecules^16^. The newly emerged RBD3 and 4 domains suggest potential evolutionary novel functions of NCL in these higher organisms. However, in contrast to RBDs 1 and 2, RBDs 3 and 4 have remained overlooked and understudied.

NCL regulates gene expression by binding both coding (mRNA) and non-coding RNA species (rRNA, miRNA, and lnc RNA). It is well established that NCL interacts with RNA preferentially through stem loop structures including apical loops or hairpin loops^20, 26^ and AU^18, 27^/G rich elements^28, 29^, both serving as signature sequence or structural motifs for NCL-RNA affinity. In fact, a G-rich stem-loop structure called nucleolin recognition element (NRE), found in pre-ribosomal RNA, establishes a primary role of NCL in processing rRNA. Similarly, NCL also demonstrates high affinity for the 11 nt single stranded evolutionary conserved motif (ECM) found 5 nt downstream of the pre-rRNA processing site^30^. Additional RNA recognition motifs which NCL is known to interact with include AU rich elements (ARE), G-quadruplex structures in the tumor suppressor *TP53* mRNA^25^ and a stem loop forming GCCCGG motif in *GADD45* mRNA in DNA damage response^29^. NCL-mRNA interactions mediated by its RBDs influence mRNA turnover rate or translation^25, 26^, while NCL-lnc-RNA binding also has implications in RNA localization^31, 32^. Overall, it is clear that NCL-RNA interactions have a profound influence on many cellular processes that control growth, proliferation, and survival.

As a member of the short non-coding RNA molecules, microRNAs (miRNA) are often dysregulated in many cancers where the aberrant miRNA processing is linked to tumorigenesis^33^. Processing of primary-miRNA (pri-miRNA) to precursor-miRNA (pre-miRNA) in animals is mediated via the microprocessor complex (MPC) in the nucleus. The pre-miRNA is then transported to the cytoplasm by shuttle proteins and subsequently processed into its mature form by Dicer^34, 35^. In plants, on the other hand, both the pri- and pre-miRNA are processed solely in the nucleus by Dicer-Like1 protein (DCL-1) and a few more helper proteins^36, 37^. A similar mechanism also exists in *Dictyostelium Discodeium*, a slime mold species, where the double stranded RNA (dsRNA) binding protein RbdB processes pre-miRNA molecules^38^. NCL is known to interact with the active components, Drosha and DCGR8 in the microprocessor complex^21^ and the emergence of NCL-RBD3-4 in higher organisms coincides with the roles of NCL in miRNA processing. We, therefore, propose that the emergence of NCL-RBD3-4 in higher organisms coincides with the NCL role in miRNA processing in evolution and that RBD3-4 possess sequence/structural determinants that specifically recognize miRNA precursor molecules in NCL protein.

The focus of this study is to elucidate the selective preference of specific NCL RBDs for the recognition of miRNAs using an *in silico* approach. Structural information for NCL RBDs and miRNA molecules is either unavailable or limited to partial structures. To fill these structural gaps, in this study we generated 3D models of the human NCL central region containing all 4 RBDs as well and various tandem pairs of RBD, as well as selected miRNAs . Our data include much needed structural models of NCL-RBDs, miRNAs and predicted scenarios of NCL- miRNA interactions from RNA-Protein docking algorithms. Our study suggests a predominant role of NCL RBDs 3 and 4 in miRNA target specificity and provides details about key motifs/residues at the NCL-substrate interface responsible of specific NCL-miRNA interactions. Structural modeling and *in silico* analysis tools provide valuable information to fill in the knowledge gaps and provide a cost effective and rational entry point in experimental design. Ultimately, the insights from this study can lead to future studies for identifying new drug design targets to regulate NCL functions in gene expression during tumorigenesis.

## Results

### 3D structural models of NCL RBD1-4, RBD3-4 and specific miRNAs provide a complete structural picture needed to study NCL-miRNA interactions

The tandem pair of NCL RBD1-2 structure is experimentally resolved (PDB ID: 2KRR)^39^ and well characterized for its interactions with various rRNA^16^ and mRNAs^25, 26^. While individual crystal structures of RBD3 and RBD4 are available, albeit misannotated (PDB IDs: 2FC9 & 2FC8^40^), the functional role(s) of the NCL RBD3-4 tandem pair remain unresolved and unexplored. Structural information for all the RBDs in NCL are critical to fully understand the mechanistic details about various RNA targets of NCL and the key features that render target-specificity. Lack of structural details on RBD1-4 as a whole and the RBD3-4 tandem pair, pose limitations in dissecting the specificity with which NCL binds to a wide-range of RNA species and plays pivotal role/s in pathophysiology. Therefore, structural models of NCL RBDs and miRNA molecules generated using template based and *ab initio* approaches were analyzed to identify robust models with high evaluation profiles **(Supporting Tables S1 and S2**) to bridge the gap in NCL RBD structural information. Figure 1 shows that the superposition of the top ranked RBD1-4 (**Figure 1A**) and RBD3-4 (**Figure 1B**) models with the existing individual crystal structures of RBD1, 2, 3 and 4 overlays very well with minor deviations in the loop regions.

**Figure 1.**
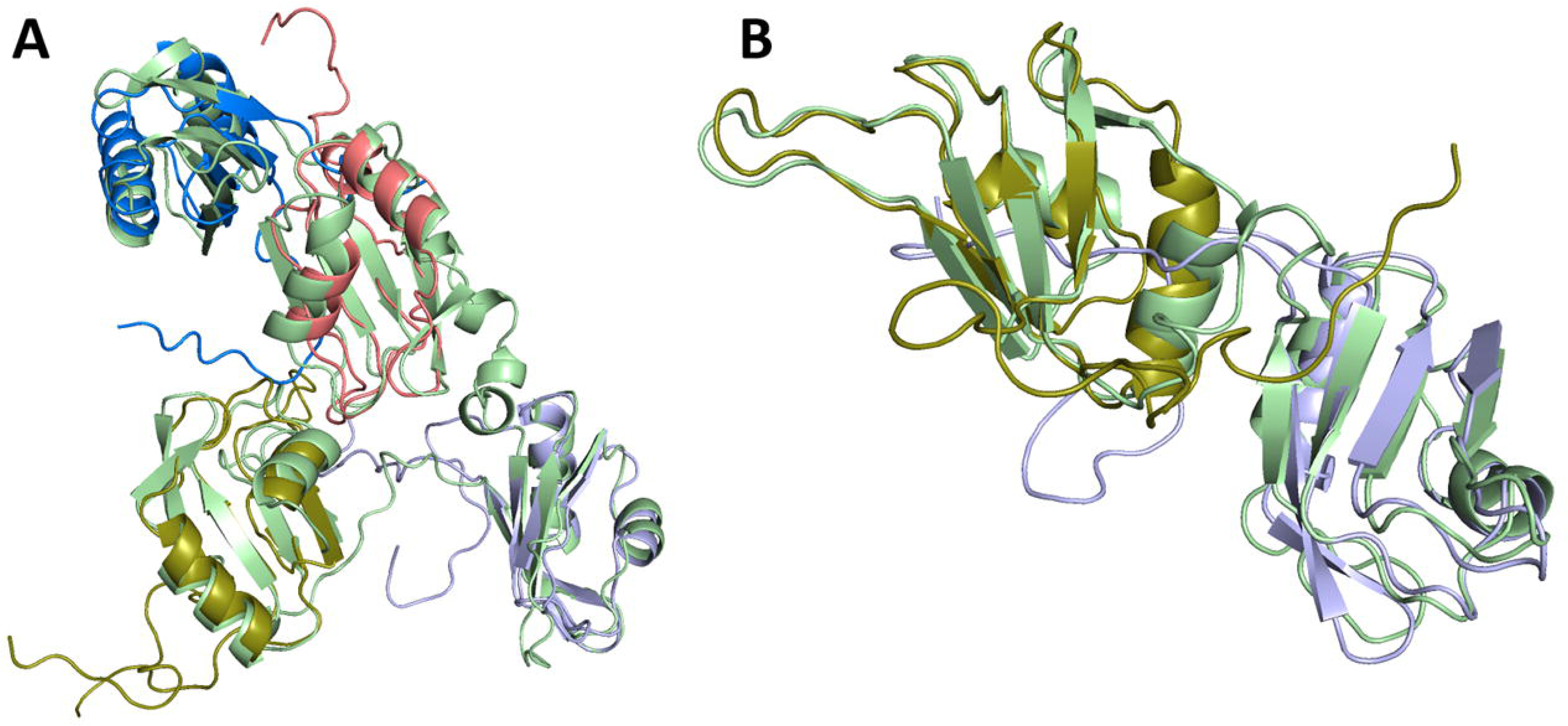
NCL tandem RBD models display high structural similarities with existing individual NCL RBD crystal structures. **A)** Superposition of the modeled RBD1-4 (pale green) with individual crystal structures of NCL RBD1 (PDB ID: 1FJ7 ;marine blue), RBD2 (PDB ID: 1FJC deep salmon), RBD3 (PDB ID: 2FC9; Lightblue), and RBD4 (PDB ID: 2FC8; deep olive). RMSD scores for RBD1, RBD2, RBD3, and RBD4 are 3.594 _, 2.324 □, 1.775 □, and 2.442 □, respectively. **B)** Superposition of the modeled RBD3-4 (pale green) with individual structures of NCL RBD3 (PDB ID: 2FC9; Lightblue), and RBD4 (PDB ID: 2FC8; deep olive) RMSD scores for RBD3 and RBD4 are 0.956 □ and 1.753 □, respectively

Previous biochemistry studies have shown that NCL interacts with pri-mir-15a, pri-mir-16-1, pri- mir103a, pri-mir-21, pri-mir-221, and pri-mir-222^21, 41^. Structural information of these miRNA molecules is largely unavailable in the current databases. Therefore, 3D miRNA models generated based on secondary structure prediction using 3D modeling programs were evaluated to identify top ranked models (**Figure 2**). Additionally, if multiple models were ranked the same in structural evaluation profiles (**Supporting Table S4**), they were all used in RNA-Protein docking analysis to check for reproducibility of results. Our predicted models provide a complete picture of the structural information for both NCL and the miRNA analyzed in this study.

**Figure 2.**
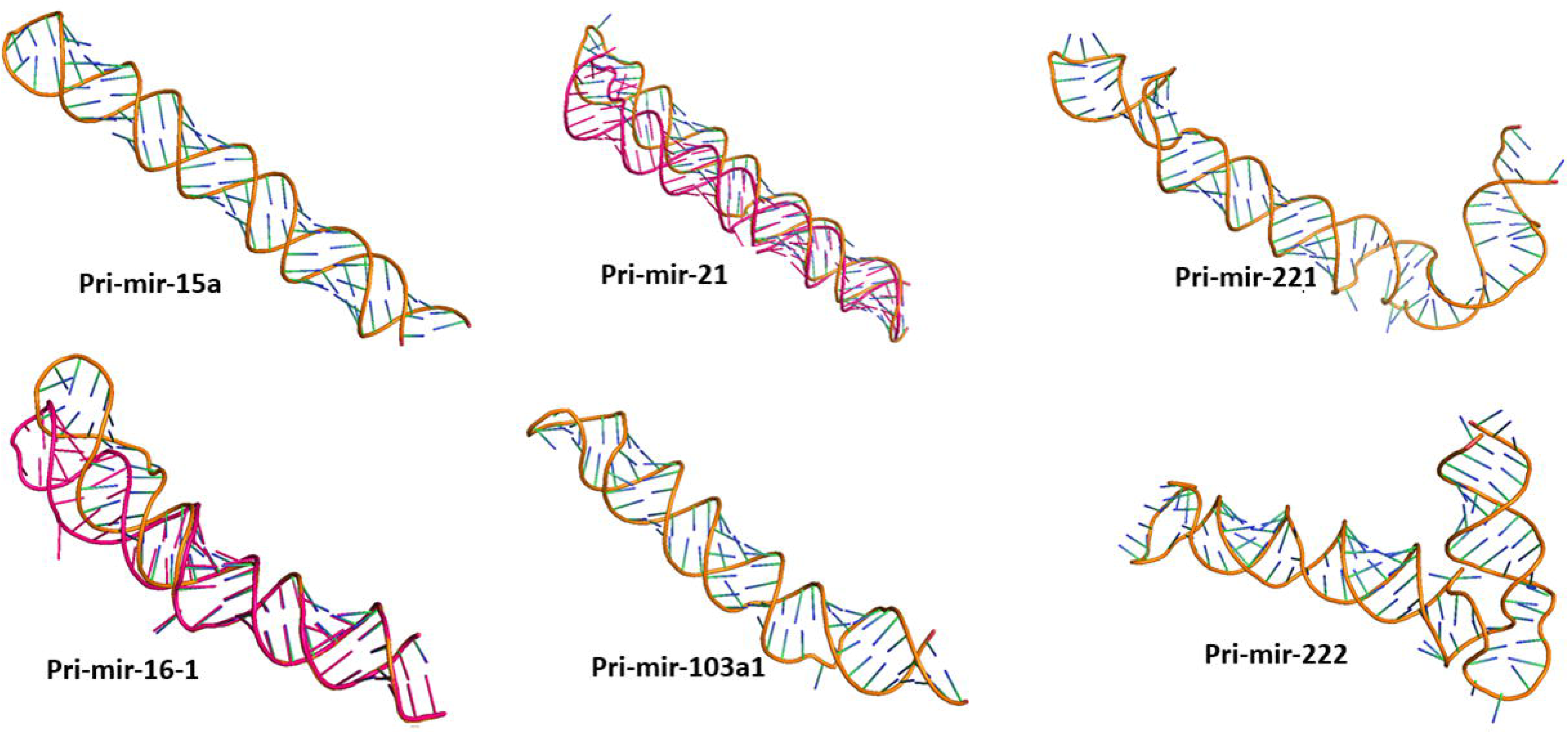
Structural models of the miRNAs analyzed in this study. Top ranked models for all 6 pri-miRNA (orange backbone with bases shown in blue). Alternative miRNA models that exhibited comparative evaluation profiles are shown in dark red.

### Docking analysis of miRNA with NCL RBDs reveals 3 possible NCL RBD-miRNA interaction scenarios

Our comprehensive analyses of NCL RBDs and a set of NCL-interacting miRNAs suggests 3 possible binding modes between NCL RBDs and miRNA molecules:

#### Mode 1: Both RBD3 and RBD4 form the RNA-protein interaction interface

In mode 1 scenario, the interface is built by the contribution of RBD3 with conserved residues on the RNP1 motif (K217, Y219, F221) as well as new interacting residues identified in RBD4 (positively charged residues on β-2 strand (R291, R293)) and on the linker loop between β-2 and β-3 (R298). This binding mode is predicted for the interactions of mir-16, mir-221, and mir-222 with NCL. In this putative binding mode, RBD3 and RBD4 interact with the double stranded RNA from opposite ends and cooperate to hold the miRNA molecule in a clasped orientation (**Figure 3**).

**Figure 3.**
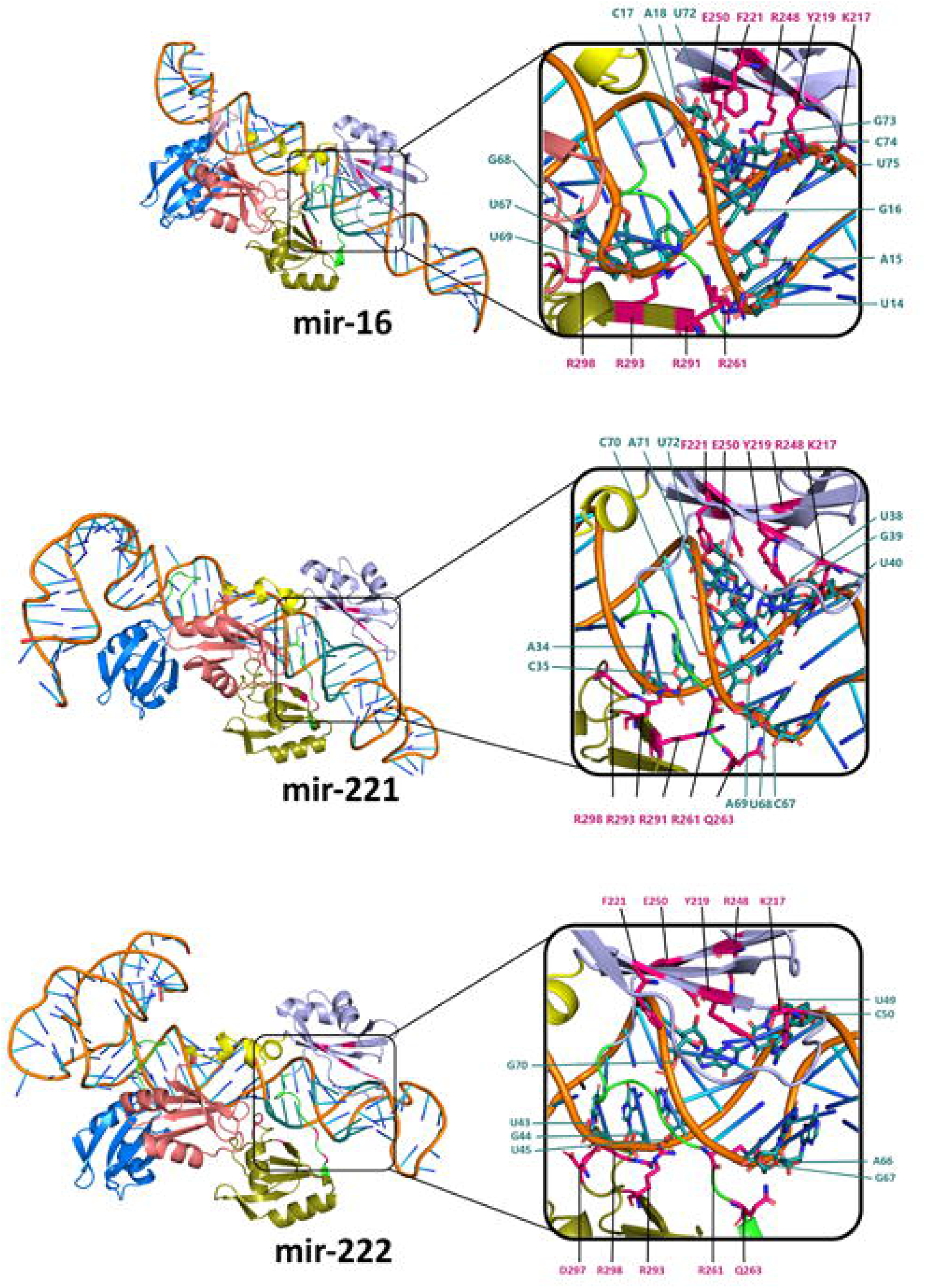
Representative docking poses exhibiting binding mode 1 involving RBD3-4. **A)** Complete docking scenarios. **B)** Zoomed-in insets with intermolecular distances between residues indicated. miRNA molecule backbone (orange), RBD1 (marine blue), RBD2 (deep salmon), RBD3 (light blue), RBD4 (deep olive). The linker regions between RBDs 1 and 2, 3 and 4 are colored in green while the linker between RBDs 2 and 3 are colored in yellow. Interacting nucleotides on the miRNA and NCL-RBDs are indicated with deep teal and hot pink, respectively.

#### Mode 2: RBD4 forms the major component of the interaction interface

This binding mode 2 features anticipated residues on RNP1 motif (K304, F306, F308), the positively charged residues on β-2 strand (R291, R293) of RBD 4, and on the linker loop between β-2 and β-3 (R298). This binding mode is predicted in the mir-15a and mir-103a docking scenarios (**Figure 4**) and suggests the involvement of multiple beta strands from a single RBD, RBD4. The binding mode 2 bears resemblance to the mir-18 binding mode of hnRNP A19 (described in the next section), both in terms of the composition and location of the residues involved in the interaction and in the overall arrangement of the beta strands. In both cases, the interactions are driven by residues on 3 beta strands of the same RBD.

**Figure 4.**
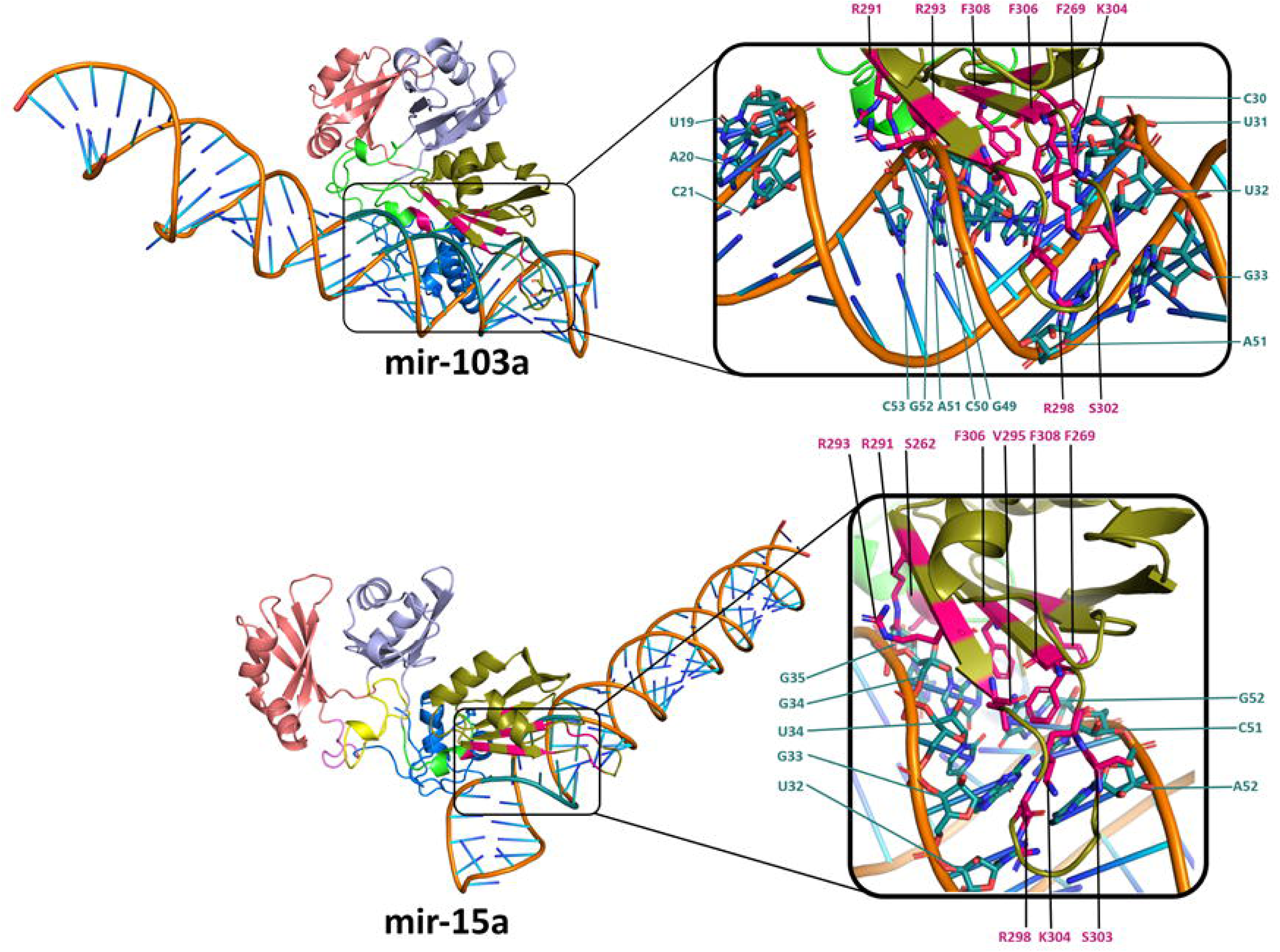
Representative docking poses exhibiting binding mode 2 involving RBD4. **A)** Complete docking scenarios. **B)** Zoomed in version with intermolecular distances between residues indicated. RBD1 (marine blue), RBD2 (deep salmon), RBD3 (light blue), RBD4 (deep olive The linker regions between RBDs 1 and 2, 3 and 4 are colored in green while the linker between RBDs 2 and 3 are colored in yellow. Interacting nucleotides on the miRNA and NCL- RBDs are indicated with deep teal and hot pink, respectively.

#### Mode 3: RBD3-4 predominantly drive the interactions with a minor contribution from RBD1-2

In this mode 3, besides the involvement of both RBD3-4 as predicted in binding mode 1, RBD1- 2 is predicted to have a minor contribution by forming salt bridges with the phosphate backbone. This binding mode (**Figure 5**) is only predicted with mir-21 docking scenarios.

**Figure 5.**
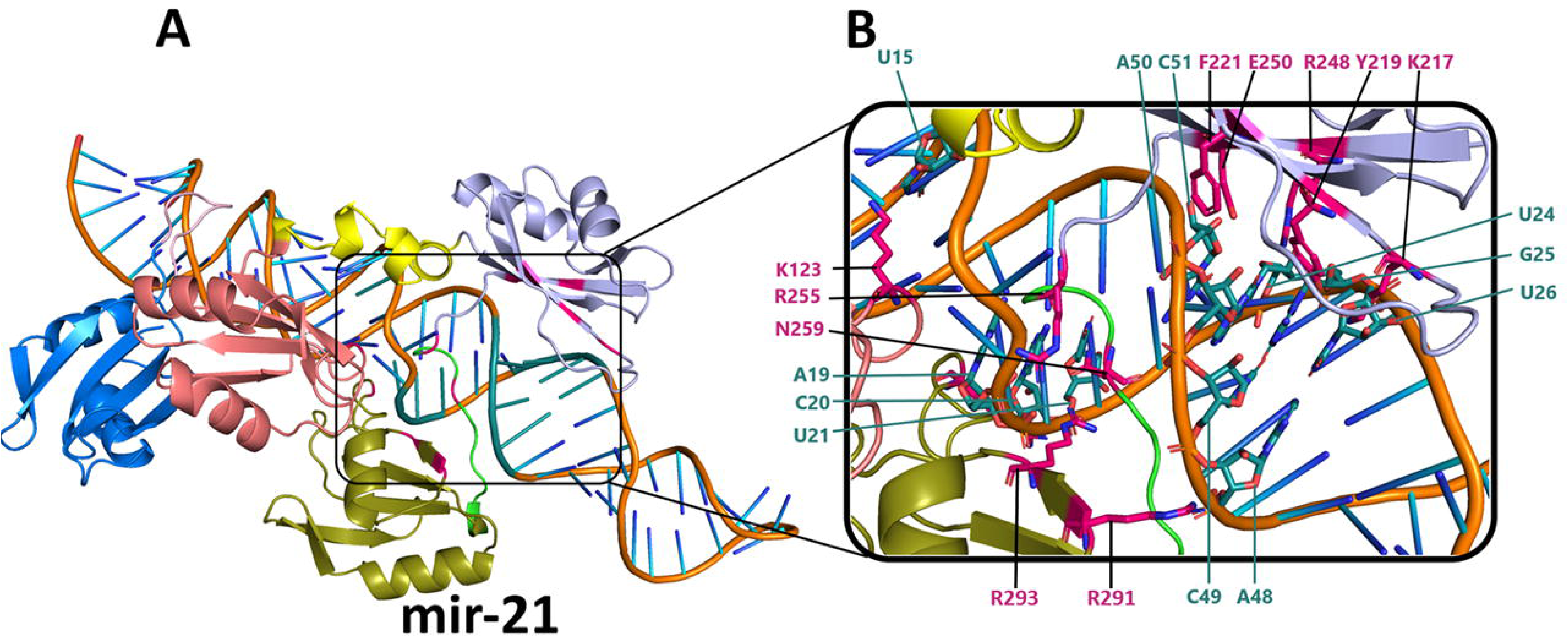
Representative docking poses exhibiting binding mode 3 involving RBD234. **A)** Complete docking scenarios. **B)** Zoomed in version with intermolecular distances between residues indicated. RBD1 (marine blue), RBD2 (deep salmon), RBD3 (light blue), RBD4 (deep olive). The linker regions between RBDs 1 and 2, 3 and 4 are colored in green while the linker between RBDs 2 and 3 are colored in yellow. Interacting nucleotides on the miRNA and NCL- RBDs are indicated with deep teal and hot pink, respectively.

### RNP motifs are conserved on all 4 NCL RBDs but RBD-miRNA interactions are driven primarily by RBD 3 and 4

An alignment of NCL RBDs with the hnRNP A1 shows the conservation of the RNP motifs as well as the key residues that are involved in hnRNP A1-RNA/DNA interactions in all 4 NCL RBDs (**Figure 6A**). However, our docking studies predict the RNP motifs of RBD3-4 are predominantly involved in miRNA interactions (see docking results, Figures 4-6). The aromatic and basic residues of RNP1 (K217, Y219, F221) of RBD3, RNP1 (K304, F306, F308) and RNP2 (F269) of RBD4 were the most frequently predicted RBD residues at the RNA-protein interface (**Figure 6B**). Additionally, the linker region connecting RBD3 to RBD4 contains several residues frequently predicted to be involved in NCL-miRNA interactions (R255, N259, R261, Q263; not shown in the figure).

**Figure 6.**
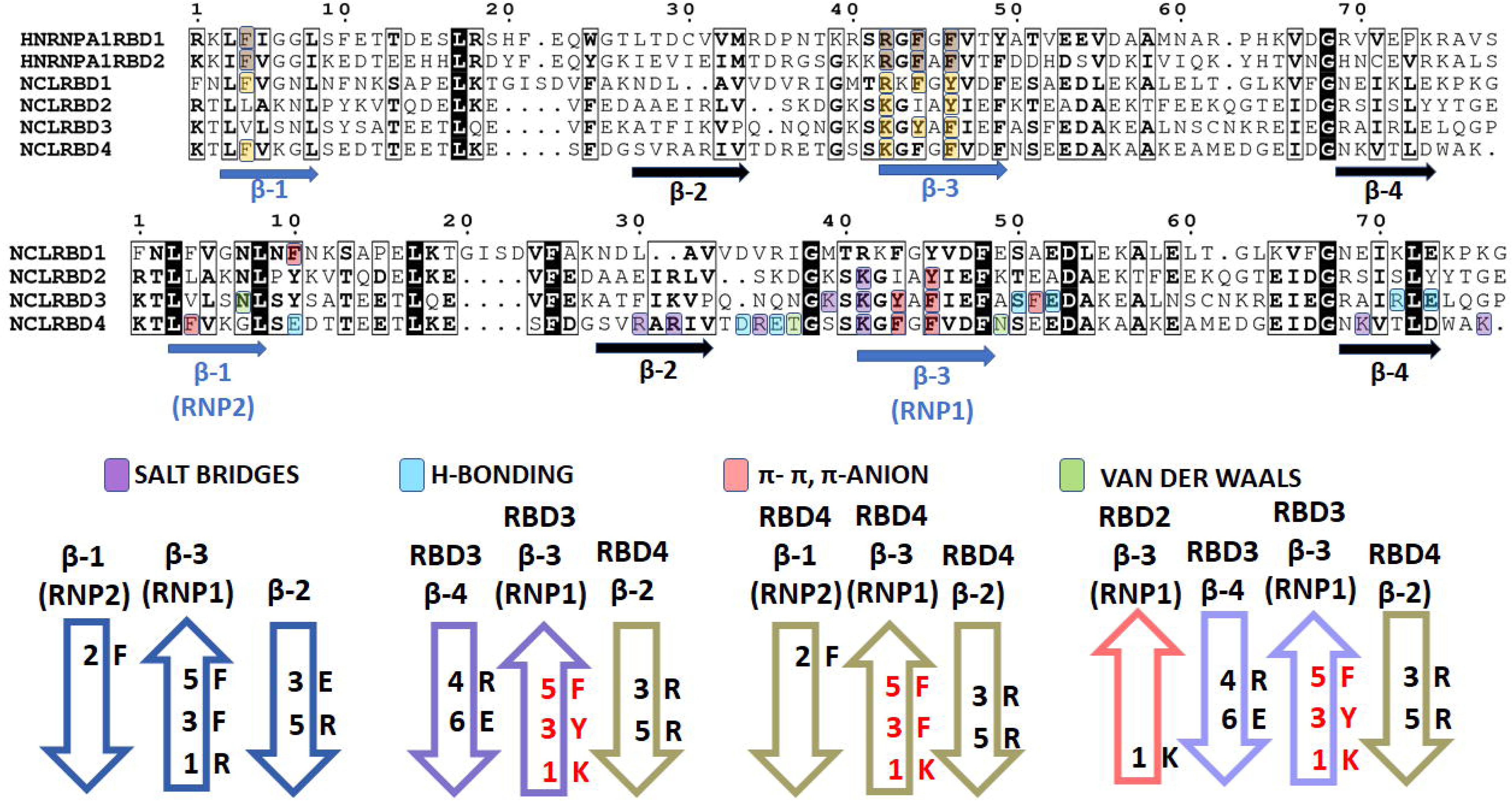
NCL residues predicted to interact with miRNAs. **A) Comparison of hnRNP A1 RBDs with NCL RBDs** (PDB ID: 6DCL). Beta strands and RNP motifs are indicated with arrows. Known hnRNP A1 residues that interact with ssDNA and miR-18 are indicated in light brown background. NCL residues conserved in the equivalent positions are indicated in yellow background. Conserved residues are indicated in bold and black background. **B) Summary of NCL-RBD residues most frequently predicted to interact with miRNA based on docking**. Different types of interactions are indicated with background colors **C) Summary of predicted NCL-miRNA binding modes and a comparison with hnRNP A1-miRNA binding model. 1)** Known hnRNP A1-mir18 interaction residues. **2)** Proposed NCL-miRNA binding mode 1 where both RBD3&RBD4 are involved in the interactions. **3)** Proposed NCL-miRNA binding mode 2 where RBD4 alone interacts with miRNA **4)** Proposed NCL-miRNA binding mode 3 which RBD3&4 and a single residue from RBD2. Beta sheets are numbered on top, and residues predicted to interact with miRNA are indicated in red. Newly identified residues are indicated in black. The numbers indicate the position of these residues in the corresponding beta strands.

These results are corroborated by a predictive analysis based solely on primary structure using the Catrapid algorithm that similarly suggests RBD3, RBD4, and the linker region between RBD3-4 have the highest RNA binding propensity for all tested miRNA molecules (**Supporting Figure S2**). Additionally, we performed a control docking of NCL RBD1-4 with miR-155, a microRNA reported to be not affected by NCL^41^. Interestingly, this docking yields noisy and inconsistent results with no clear selectivity for any particular RBD (**Supporting Figure S3**), supporting the quality of our *in silico* prediction. Similarly, results of the docking experiments with mutant RBD3-4 models revealed that RNP motifs on RBD3 and 4 lose the specific recognition of miRNA interactions observed in wild type NCL.

### Aromatic and basic residues play a key role at the NCL RBD3-4-miRNA interface

RNA-NCL docking scenarios of both RBD3-4 and RBD1-4 with miRNA models highlight the role of some of the positively charged arginine residues in β-2 (R291, R293) as well as the linker between β-2 and β-3 on RBD4 (R298) as these residues are consistently predicted to be involved in nearly all of the docking scenarios analyzed. These basic residues form salt bridges with the phosphate backbone of the various miRNAs. Similarly, our results indicate that aromatic residues on β-1 (F269 in RBD4) and β-3 (Y219 and F221 in RBD3 & F306 and F308 in RBD4) interact with miRNA molecules through π- π stacking and π-anion interactions with the nucleobases and the phosphate backbone, respectively. Importantly, the docking analysis of RBD1-4 models align with those of RBD3-4 models (**Supporting Figure S4**). Our results also show that residues on RNP motifs from RBD1-2 are largely not involved in NCL RBD-miRNA interactions, despite the similar functional profile of the RNP motifs. A control docking analysis using only RBD1-2 yields noisy scenarios with no clear trend (**Supporting Figure S5**), again reinforcing the idea that NCL RBD3-4 preferentially interact with the miRNA under study. Based on the docking results, we show that NCL-RBDs interact with pri-miRNA nucleotides almost exclusively within the region that corressponds to the mature miRNA molecule. In our results, NCL is predicted to interact frequently with G-U pairs and kink turns (Figure 7). Interactions were observed predominantly at the major groove regions of the double stranded miRNA molecules as these regions were more accessible when compared to the minor groove regions.

**Figure 7.**
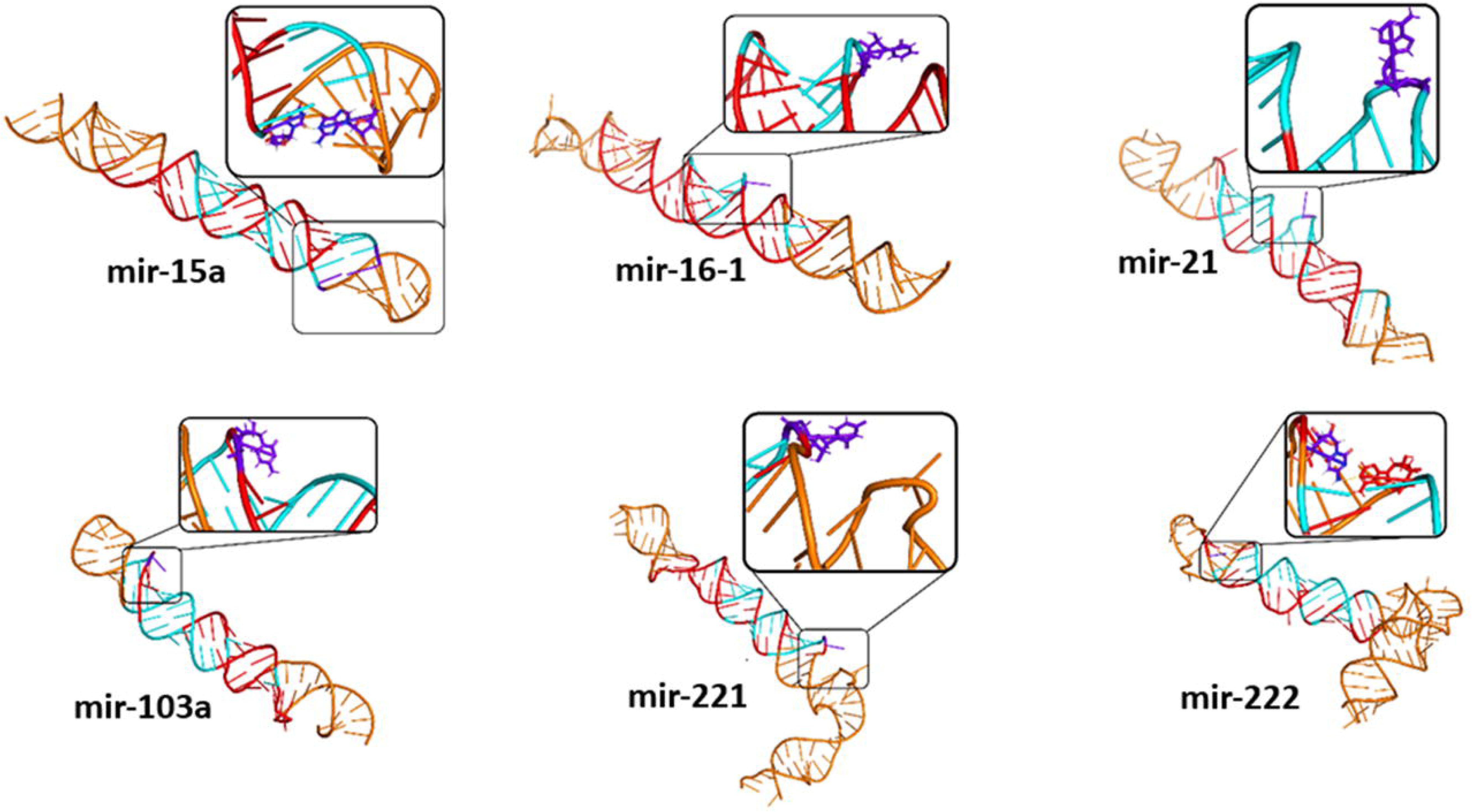
Location of non-canonical base pairing nucleotides and motifs in miRNA that interact with NCL-RBDs. miRNA residues most frequently predicted to interact with NCL- RBDs are show in cyan for all miRNA structures where pri-miRNA is shown in orange and region of the mature miRNA is shown in red. Non-canonical base pairs and kink-turns are indicated in dark purple.

## Discussion

The importance and functionality of RNP motifs in RBP-RNA interactions are well established in previous studies^9^. Since there is a high sequence similarity between NCL and hnRNP A1 RBDs, and hnRNP A1-RNA interactions are known, a preliminary prediction of putative residues that may be important in NCL-miRNA interactions could be made based on sequence conservation between NCL and hnRNP A1. The RBP-hnRNP A1 interacts with MPC and pri-mir-18 by utilizing aromatic/charged residues found on its RNP motifs as well as on non-RNP beta strands^9^. These aromatic/charged residues show strong conservation between NCL and hnRNP A1 sequences suggesting existence of a similar RNP-mediated mechanism in NCL-RNA interactions (**Figure 6A**). This study confirms this prediction through detailed docking analyses *in silico*. Our results provide three alternative NCL-miRNA binding possibilities with common underlying conserved residues at an equivalent position as in hnRNP A1-RBD taking part in RNA binding (**Figure 6C**). Aromatic residues in the RNP motifs are known to be capable of initiating interactions with multiple nucleobases at the same time through stacking interactions and promoting structural stability to the RNA-RNP binding motif ^10^. We consistently predicted the involvement of these aromatic residues in all binding modes of miRNA-NCL. Additionally, we frequently predicted some charged residues on different beta strands from the same RBDs to be involved in NCL-miRNA interactions. Several arginine residues (R291, R293, and R298) were consistently predicted to interact with phosphate groups on the nucleotides via salt bridges (e.g., R293 with G16 in Fig 3, and with A34, and U45 in the figures for mir-16, mir-221, and mir-222, respectively). These residues provide additional structural stability to RNA-protein interactions similar to previous studies^9^. Our docking results also revealed that each NCL-RBD interacts with the miRNA duplex structure from opposite sides and forms a clasp around it. Linker regions between RBD3 and RBD4 were consistently predicted in several scenarios to hold the miRNA molecule from an additional third side, thus tightening the NCL-RBDs grip on miRNA (**Figures 4-6**).

Our results indicate that NCL RBDs demonstrate preference or affinity for interaction with certain types of miRNA motifs. It was previously established that NCL prefers interacting with RNA loops while driving ribosomal biogenesis^42^. Non-canonical base pairing is known to lead to kink-turns, bulges, mismatched pairs, and wobble pairs in miRNA structures and presents suitable interaction sites that would be unavailable otherwise. In canonical pairs such as G-C or A-U, amino groups of each base-pair are projected into the major groove, creating a region with positive electrostatic potential^43^. The G-U base-pair is an example of non-canonical base pair, where oxygen groups from both nucleotides face the major groove side instead of the amino groups and leading to negative electrostatic potential in the region of the major groove of the dsRNA^43^. Our results also highlighted the preference of NCL RBDs on non-canonical base pairs when interacting with miRNAs (**Figure 7**). Such regions are expected to interact with amino acids with positively charged side groups such as arginine or lysine^43^. This ties in with our results as several arginine and lysine residues from NCL RBD3-4 were predicted as some of the most frequently encountered residues in the various docking scenarios obtained.

Additionally, wobble pairs and mismatched pairs are known to be important elements of primary miRNA processing by the MPC^44^. Certain RNA motifs such as UGU/GUG are known to be enriched around the apical loop regions^45^ and the preference of NCL to interact with regions close to the apical loops has been demonstrated in previous studies investigating NCL-rRNA^42^ and NCL-mRNA^26^ interactions. This consensus sequence is also important in pri-miRNA processing by the MPC as DGCR8 is thought to recognize and interact with this consensus sequence^46^. Our results revealed that NCL RBDs were able to recognize this UGU/GUG motif for all miRNA molecules tested in this study (**Figure 8**). In the cases of mir-15a and mir-103a, these regions were identified adjacent to apical loop structures. It is also interesting to note that, in binding mode 2 (observed in mir15a and mir103a), RBD4 by itself seems sufficient to drive NCL-miRNA interactions (**Figure 8A**). In the cases of mir-21 and mir-16-1, NCL interacts with regions distant from apical loops, but closer to bulge regions towards the middle of the miRNA molecule. (**Figure 8B**). Since DGCR8 also recognizes the same motif for interactions, our results suggest that NCL potentially binds adjacent to DGCR8 or could even replace it in as a possible scenario of miRNA – NCL interactions. Alternatively, in the cases of mir-221 and mir-222, NCL-miRNA interactions were predicted to be localized slightly distant to the apical loop region and closer to the basal stem region containing a mismatched GHG/CUC motif (**Figure 8C**). This manifests as a mismatched bulge, which is a common element in most pri-miRNA structures^46^. Drosha cleaves its miRNA targets around the basal stem close to this motif^46^. This cooperation model suggests that NCL RBDs binds closer to Drosha. In both scenarios, these motifs likely serve as anchoring points for NCL RBDs when they interact with miRNA in cooperation with MPC.

**Figure 8.**
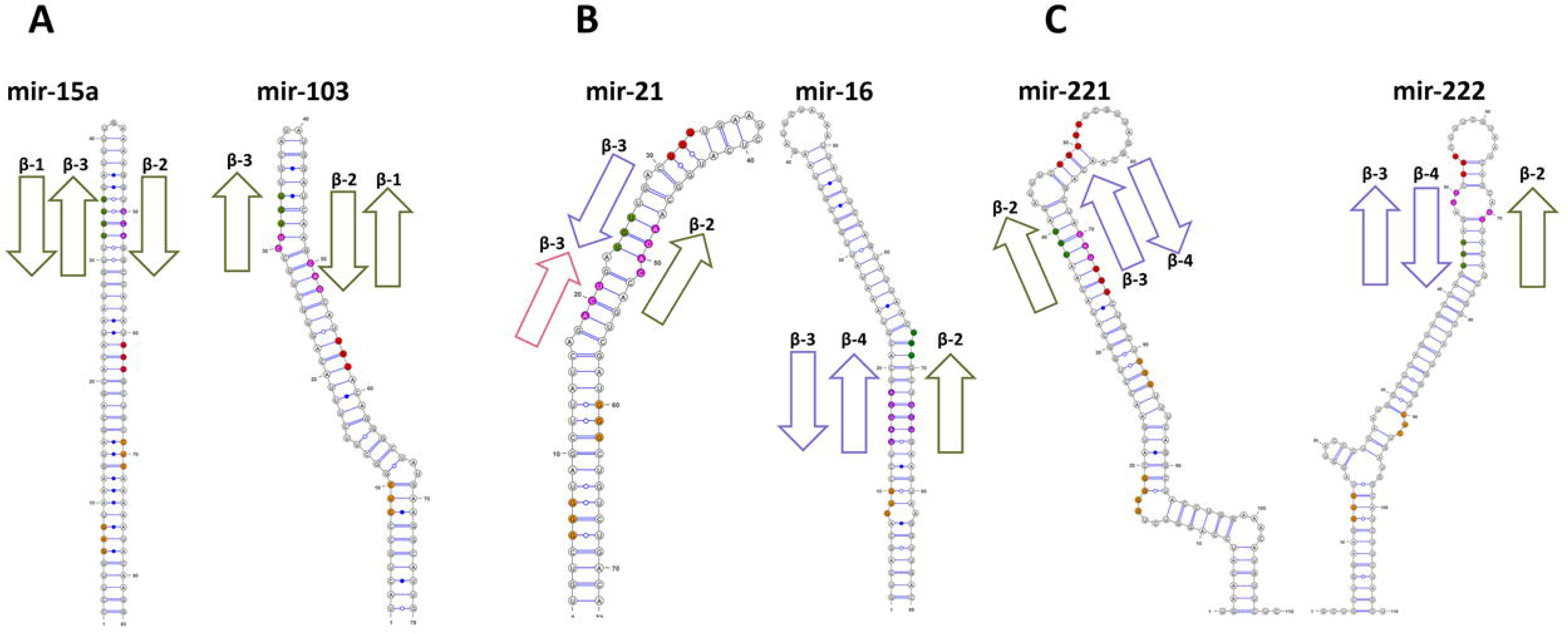
Mapping of miRNA residues interacting with NCL. **A)** NCL contacts miRNA molecules at the UGU/GUG motifs closest to the loop. **B)** NCL contacts miRNA molecules on UGU/GUG motifs distant from the loop and closer to GHG/CUC motifs. **C)** NCL contacts longer miRNA molecules on UGU/GUG motifs slightly distant from the apical loop and closer to the loop stem. Beta strands are numbered on top of the arrowheads. RBD1 (marine blue), RBD2 (deep salmon), RBD3 (light blue), RBD4 (deep olive), and linker loops between RBDs (neon green). UGU/GUG motifs on miRNA that NCL interacts are indicated in green. miRNA residues that NCL interacts with are but is not part of an UGG/GUG motif is indicated in purple. UGU/GUG motifs on miRNA that NCL is not interacting are indicated in red. GHG/CUC motifs are indicated in orange.

Similarly, multiple other RBPs including hnRNP A1, Lin28B, RBFOX3, and HuR interact with either the terminal loop or the stem regions of the pri-miRNA molecules to either promote or suppress pri-miRNA processing by the MPC proteins depending on the location of the interactions and the targeted miRNA^47^. For example, both Lin28B and hnRNP A1 interact with the terminal loop of a subset of pri-miRNA structures. hnRNP A1 is predicted to help with the processing of pri-miR18-a by binding to the terminal loop and causing a relaxation of the miRNA structure and therefore making it easier for MPC to interact with miRNA^48^. However, when hnRNP A1 interacts with let-7 pri-miRNA terminal loop, it outcompetes binding of another RBP, KH-type slicing regulatory protein (KSRP) known to promote biogenesis of let-7^48^ and decreases pri-miRNA processing by Drosha. Lin28B, on the other hand, always negatively regulates this process by interacting with the terminal loop and inhibiting Drosha from interacting with miRNA^49^. Both RBFOX3 and HuR bind to the basal stem and inhibit miRNA processing by blocking catalytic activity of Drosha^50, 51^. Based on these studies, it is clear that the relationship of RBPs with the MPC is both location and context dependent.

A recent structural study investigating interactions of the MPC with pri-miR16-2 revealed that 2 DGCR8 proteins interact with nucleotides adjacent to the apical loop region of the pri-miRNA molecule^52^. Canonical function of DGCR8 is to interact with pri-miRNA using its double stranded RNA binding domains (dsRBDs) and present the pri-miRNA molecules to Drosha for processing^52^. This is illustrated in **Figure 9A**. Many RBPs can interact with regions of pri-miRNA molecules that are also regions where DGCR8 proteins bind. We speculate that NCL- RBDs promotes pri-miRNA processing of certain miRNAs by a similar mechanism as observed in hnRNP A1-mir18 interactions. As a RBP capable of binding double stranded RNA molecules, NCL could potentially replace the pri-miRNA presenting functions of one or both DGCR8 proteins. Our results suggest that for certain miRNA, NCL can wrap around the double stranded miRNA molecule with two RBDs like DGCR8 (**Figure 9B**). Since the lengths of oncogenic miRNA transcripts are variable, we speculate that NCL may act as a bridging agent between DGCR8 and Drosha when processing longer transcripts (**Figure 9C**). Our findings present a snapshot of NCL-miRNA that give an initial insight into these interactions. We envision future studies to elaborate the NCL-miRNA interaction dynamics over longer time-scales to get a finer grained picture of the underlying molecular mechanisms and their downstream effects.

**Figure 9.**
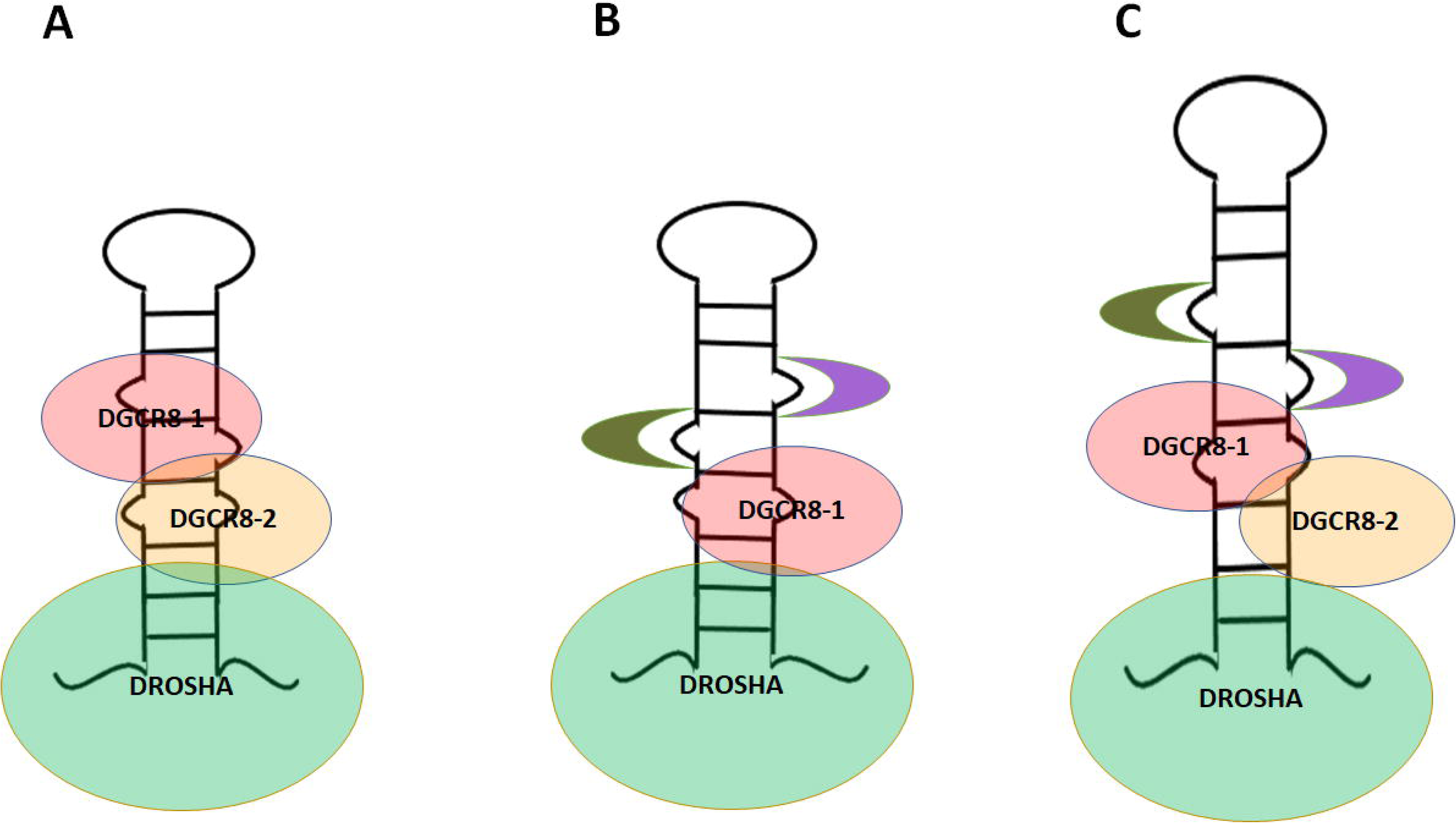
Canonical and proposed models of NCL, Drosha, and DGCR8 interactions with pri-miRNA molecules. **A)** Canonical model involving Drosha & DGCR8-1 and 2^52^ **B)** Hypothetical model A where NCL-RBDs could replace DGCR8 – 2 **C)** Hypothetical model B where all MPC proteins and either one or both NCL-RBDs interact with the pri-miRNA molecule. NCL-RBD3&4 are colored in light blue and deep olive, respectively. DROSHA is light green. DGCR8-1 is red, and DGCR8-2 is orange.

Drosha and DGCR8 interact with different RNA types including precursor-mRNA (pre-mRNA) and non-coding RNA and are also involved in double stranded DNA break repair mechanisms^53^. Since NCL and MPC proteins can interact in a non-miRNA context^21^, it is quite likely that they cooperate in other biological pathways as well. Studies testing NCL- MPC cooperation in pre-mRNA processing may be a natural extension of this study since NCL is also known to interact with mRNA UTR molecules to manipulate their expression levels in human cancers^20, 25, 26^.

NCL cellular localization is highly complex and context-dependent. Pathophysiological functional implications of NCL are even more enigmatic. Despite this multidimensional behavior, NCL is a promising target for cancer therapeutics. Therapeutics such as the immunoagents 4LB5-HP-RNase^54^, G-rich DNA oligonucleotide, aptamer AS1411^55^, and antagonist pseudopeptides, N6L^56^ and HB19^57^, target either surface or cytoplasmic NCL to regulate its functions in miRNA synthesis, RNA metabolism, cell proliferation, angiogenesis and metastasis, in a variety of cancer types including breast cancer.

In summary, we predict two putative binding modes where NCL RBD3-4 specifically drive NCL-miRNA interactions and an alternative binding mode where a small contribution from RBD12 is also involved. As we had hypothesized RNP motifs on RBD3-4 play a significant role in these interactions; additionally, we report novel residues important for the interaction that have not been identified in any other previous study. Importantly, our data support an idea that the exclusive presence of NCL RBD3-4 in animals might be evolutionarily relevant for NCL functions as drivers for microRNA processing. Our data delineate critical residues in NCL and recognition motifs in the miRNA important for NCL-miRNA interactions. Future studies designed to validate critical residues in these motifs that are important for interactions, using site-directed mutagenesis in cellular conditions will be required. Once confirmed experimentally, NCL-miRNA interacting interface provide a valuable drug targets for the development of cancer therapies to control specific gene expression.

## Materials and methods

A flowchart summarizing the general protocol of the present study with the various tools used to analyze NCL-miRNA interactions (discussed in detail below) is shown in the supporting Figure S1.

### Sequence analysis and generation of 3D models of NCL-RBDs

Since only partial structural information for NCL-RBDs is available, robust 3D models of all 4 NCL RBDs (RBD1-4), and RBD3-4 were built; structural information for RBD1-2 in tandem (PDB ID: 2KRR) ^39^ and for individual RBDs (PDB ID: 1FJ7 & 1FJC, respectively)^42^ is available for human NCL. The human NCL sequence available from NCBI database ^58^ was analyzed for its domain architecture using the programs SMART^59^, Pfam^60^, Uniprot^61^, and Interpro^62^ to confirm the domain boundaries of the individual RBDs accurately as well as to identify any potential sequence motifs of relevance. The multiple sequence alignment tool Clustal Omega^63^ was used to align the NCL-RBDs with hnRNPA1 RBDs to identify conserved residues. The multiple sequence alignment was visualized using the alignment editor Espript3^64^.

Delineated tandem domain pairs were modeled using both template-based methods (Swissmodel^65^ Intfold^66^, Phyre2^67^ and *ab initio* modeling approaches (Robetta^68^, QUARK^69^, and I-TASSER^70^) to generate structural models. To identify high quality models, the constructed models were rigorously evaluated by model verification programs including Verify3D^71^, VoroMQA^72^, Prosa-web^73^, and ProQ3^74^ (Supporting Tables S1 and S2) and correlation of their biophysical and structural properties with experimental observations. Top models were refined using ModRefiner^75^ and SCWRL4^76^ and then re-evaluated. ModRefiner first modifies the protein side chain packing by adding atoms and improves the structural quality of reconstructed models by energy minimization procedures. SCWRL4 focuses on side chain refinements to improve the models. Top scoring models were chosen for further analysis (Supporting Tables S1 and S2).

### Generation of 3D models of miRNA implicated in interactions with NCL

Structural information of miRNA molecules implicated to play a role in breast cancer (Supporting Table S3) and interact with NCL is unavailable in the databases. The limited structural data available in the protein databank for the 6 miRNA molecules under study corresponds to either partial pre-miRNA structures or apical loops (PDB ID: 2MNC, 5UZT)^77, 78^ and neither provide structural information for the complete pri-miRNA structures. Therefore, all pri-miRNA molecules were modeled using the primary sequences of miRNA stem loop structures obtained from miRbase^79^. The secondary structure of these miRNA molecules was predicted using various programs including RNAStructure^80^, RNAFold^81^, MC-Fold^82^, CentroidFold^83^, and SPOT-RNA^84^. Although all these tools provide predictions based on experimental data found in databases, some of the newer methods such as CentroidFold and SPOT-RNA also utilize machine learning in making these predictions. All of the predicted secondary structures (dot bracket format) obtained from these tools were input into the RNA modeling software including RNAComposer^85^, ifold2^86^, RNAfold3^87^, 3DRNA v2^88^ and simRNAweb^89^ to obtain 3D models of these miRNA molecules. The generated models were evaluated for their structural quality using MolProbity^90^ (Supporting Table S4). The top models were refined using RNAfitme^91^, a program that reconstructs nucleobases and nucleosides while keeping the sugar phosphate backbone fixed and aiming to reduce steric clashes.

### Prediction of motifs involved in NCL-miRNA interactions

NCL-miRNA interactions were investigated using the protein-RNA docking software HDOCK^92^ and NPDock^93^. Both programs employ a rigid body docking approach followed by scoring and clustering of structures with the lowest minimum energy followed by refinement of the results through Monte Carlo simulations. Both the programs are considered the top ranked algorithms for protein-RNA docking and score well in benchmarks. The docking results were visualized in PyMOL^94^, and the interactions mapped out using Nucplot from PDBSum^95^. The key interacting residues in the NCL-RBDs were identified based on the 10 top scoring scenarios from each docking run. These residues were compared with putative residues of NCL-RBDs identified using catRAPID^96^, a program that predicts binding propensity of a protein sequence to a given RNA sequence by measuring physicochemical properties and shape complementarity. Additionally, control docking experiments were conducted with miRNA models (mir-155) which has been shown not to interact with NCL^41^. . Similarly, docking experiments were performed using in *silico* mutagenesis at key residues of NCL identified to be important in NCL-miRNA interactions in the current study (Y219A, R261A, F311A).

## Supporting information

Supplementary Tables and Figures references

## Acknowledgements

We acknowledge the contribution of Michael Scarpati, Rahimah Ahmad, and Yue Qui for the preliminary analysis that paved the way for this work. We are extremely grateful to Dr Aaron Frank and Dr Tamar Schlick for their valuable feedback and the critical review of this manuscript. This work was partially supported by the CUNY Doctoral Students Research Grant (Round 14/2019-2020) and Provost’s pre-dissertation summer science research award (2019) to AS.

## Abbreviations

NCL: nucleolin
RBP: RNA binding protein
RBD: RNA binding domain
miRNA: microRNA
Pri-miRNA: Primary microRNA
Pre-miRNA: Precursor microRNA
mRNA: messenger RNA
RNP: ribonucleoprotein
NRE: nucleolin recognition element
ECM: evolutionary conserved motif
ARE: AU rich elements
lnc-RNA: long non-coding RNA
dsRNA: double stranded RNA
Dicer like protein-1: DCL-1
hnRNP-A1: heterogeneous nuclear ribonucleoprotein A1
MPC: microprocessor complex
DGCR8: DiGeorge Syndrome Critical Region Gene 8
Lin28B: Lin28-homolog B
RBFOX3: RNA binding protein fox-1 homolog (c. elegans) 3
HuR,: human antigen R

---

Table S1 : Evaluation and refinement of RBD14 models.

Table S2: Evaluation and refinement of RBD34 models.

Table S3: NCL interacting miRNAs based on experimental evidence or predicted interaction with NCL.

Table S4: Evaluation and refinement of miRNA models.

Figure S1: The workflow and tools used in this study.

Figure S2. CatRAPID results of NCL RBD14 for all miRNA sequences used in this study.

Figure S3. Inconsistent interactions displayed by RBD14-mir155 docking results.

Figure S4. MSA map of RBD34 predicted to be involved in NCL-miRNA interactions.

Figure S5. Inconsistent scenarios of RBD12-mir16 interactions.

