## Supplementary Table S1 for "*In silico* study predicts a key role of RNA-binding domains 3 and 4 in nucleolin-miRNA interactions"

**Table S1 : Evaluation and refinement of RBD1-4 models.**RBD1-4 models were evaluated by the structural quality control programs listed in the columns. Top scoring models from each group for every program are indicated in bold. Models which have obtained a better high score on overall were selected for refinements.

| **RBD14** | **VERIFY3D (residues with 1D-3D score above 0.2 %)** | **VoroMQA (Global score)** | **ProSA Web (Z-score)** | **ProQ3 (Global score)** | **Candidate models** | **Top ranked model** |
|---|---|---|---|---|---|---|
| **Model Evaluation** | | | | | | |
| RBD14 SWISS | | | | | | |
| 5VSU.1A | **92.63** | **0.381** | **-6.65** | **0.471** | **X** | **X** |
| 6ASO.1A | 90.86 | 0.362 | -6.31 | 0.419 | | |
| RBD14 Intfold | | | | | | |
| M1 | 62.24 | **0.403** | **-7.75** | 0.350 | **X** | |
| M2 | 57.35 | 0.392 | -7.6 | 0.360 | | |
| M3 | 65.19 | 0.387 | -7.11 | 0.449 | | |
| M4 | 80.88 | 0.314 | -4.88 | 0.510 | | |
| M5 | **88.53** | 0.312 | -5.00 | **0.517** | **X** | |
| RBD14 I-TASSER | | | | | | |
| M1 | **85.29** | **0.361** | **-6.55** | 0.408 | **X** | |
| M2 | **85.29** | 0.320 | -6.21 | 0.398 | | |
| M3 | 81.47 | 0.328 | -6.44 | **0.489** | **X** | |
| RBD14 Phyre | 73.24 | 0.294 | -5.54 | 0.421 | | |
| RBD14 Robetta | | | | | | |
| M1 | 95 | 0.414 | **-7.22** | 0.490 | **X** | |
| M2 | 99.71 | **0.423** | -7.2 | 0.486 | | |
| M3 | 99.71 | 0.385 | -7.08 | 0.341 | | |
| M4 | 92.06 | 0.401 | -7.1 | 0.480 | | |
| M5 | **100** | 0.408 | -6.79 | **0.501** | **X** | |
| **Refinement of top models** | | | | | | |
| SWISS 5VSU MR | 93.51 | 0.361 | -6.36 | 0.483 | | |
| ITASSER M1 MR | 91.47 | 0.358 | -6.31 | 0.404 | | |
| ITASSER M3 MR | 85.88 | 0.326 | -6.5 | 0.389 | | |
| Intfold M1 MR | **100** | **0.414** | **-8.13** | 0.338 | | |
| Intfold M5 MR | 86.18 | 0.374 | -5.6 | 0.509 | | |
| Robetta M1 MR | 93.24 | 0.380 | -7.15 | 0.452 | **X** | **X** |
| Robetta M5 MR | 100 | 0.386 | -6.76 | **0.486** | **X** | **X** |
| SWISS 5VSU Scwrl | 88.50 | 0.368 | -6.65 | 0.456 | | |
| ITASSER M1 Scwrl | 77.94 | 0.311 | -6.55 | 0.432 | | |
| ITASSER M3 Scwrl | 77.35 | 0.333 | -6.44 | 0.453 | | |
| Intfold M1 Scwrl | 59.88 | **0.402** | **-7.75** | 0.374 | | |
| Intfold M5 Scwrl | 79.12 | 0.332 | -5.00 | **0.506** | | |
| Swiss MR Scwrl | 92.63 | 0.364 | -6.36 | 0.479 | | |
| Robetta M1 Scwrl | 95 | 0.377 | -7.22 | 0.448 | | |

| | | | | |
|---|---|---|---|---|
| Robetta M5 Scwrl | **100** | 0.383 | -6.79 | 0.440 |
