## Supplementary Table S2 for "*In silico* study predicts a key role of RNA-binding domains 3 and 4 in nucleolin-miRNA interactions"

**Table S2: Evaluation and refinement of RBD3-4 models.** RBD3-4 models were evaluated by the structural quality control programs listed in the columns. Top scoring models from each group for every program are indicated in bold. Models which have obtained a better high score on overall were selected for refinements.

| RBD34 | MODELS | VERIFY3D (residues with 1D-3D score above 0.2 %) | VoroMQA (Global score) | ProSA Web (Z-score) | ProQ3 (Global score) | Candid ate models | Top ranked model |
|---|---|---|---|---|---|---|---|
| **Model evaluation** | | | | | | | |
| **Swissmodel** | 6DCL.1A<br>4PKD.1B | **100**<br>89.44 | **0.389**<br>0.344 | **-6.37**<br>-5.92 | **0.619**<br>0.585 | <br>X | |
| **QUARK** | M1<br>M2<br>M3<br>M4<br>M5 | 98.76<br>99.3<br>**100**<br>**100**<br>**100** | 0.341<br>0.348<br>0.360<br>0.327<br>**0.369** | -5.85<br>**-6.02**<br>-5.28<br>-5.68<br>-5.68 | 0.416<br>0.549<br>0.549<br>0.493<br>**0.569** | <br><br><br><br>X | |
| **I-TASSER** | M1<br>M2 | **100**<br>90.06 | **0.391**<br>0.371 | **-7.44**<br>-6.42 | 0.572<br>**0.623** | <br>X | |
| **IntfoldM5** | M1<br>M2<br>M3<br>M4<br>M5 | **100**<br>**100**<br>**100**<br>98.14<br>98.14 | 0.382<br>0.381<br>**0.397**<br>0.370<br>0.370 | -6.63<br>-6.83<br>**-7.04**<br>-6.45<br>-6.45 | 0.650<br>0.579<br>**0.653**<br>0.609<br>0.609 | <br><br>X<br><br> | |
| | RBD34 Phyre | 47.50 | 0.200 | -4.49 | 0.390 | | |
| **Robetta** | M1A<br>M1B<br>M2A<br>M2B<br>M3A<br>M3B<br>M4A<br>M4B<br>M5A<br>M5B | **100**<br>**100**<br>**100**<br>**100**<br>**100**<br>**100**<br>**100**<br>**100**<br>**100**<br>**100** | 0.433<br>0.437<br>0.433<br>0.428<br>0.410<br>0.420<br>0.424<br>0.424<br>**0.444**<br>0.437 | -7.1<br>-7.2<br>-7.32<br>-7.35<br>-6.9<br>-6.96<br>-7.07<br>-7<br>-7.03<br>**-7.39** | **0.724**<br>0.700<br>0.705<br>0.695<br>0.644<br>0.684<br>0.705<br>0.684<br>0.713<br>0.687 | X<br><br><br><br><br><br><br><br>X<br> | |
| **Refinement of top models** | | | | | | | |
| **Scwrl4** | Intfold<br>M3<br>ITASSER<br>M1<br>6DCL.1A<br>QUARK<br>M5 | 99.38<br>**100**<br>93.79<br>99.38<br>100<br>100 | 0.403<br>0.386<br>0.370<br>0.331<br>0.433<br>**0.444** | -7.04<br>**-7.44**<br>-6.37<br>-5.68<br>-7.1<br>-7.03 | 0.609<br>0.574<br>0.616<br>0.530<br>**0.724**<br>0.713 | <br><br><br><br><br>X | |

| | | | | | | | |
|---|---|---|---|---|---|---|---|
| | Robetta<br>M1A<br>Robetta<br>M5A | | | | | | |
| **Modrefiner** | 6DCL.1A<br>MR<br>Quark<br>m5 MR<br>Intfoldm<br>3 MR<br>I-TASSER<br>M1 MR<br>Robetta<br>M1A MR<br>Robetta<br>M5A MR | **100**<br>**100**<br>**100**<br>**100**<br>**100**<br>**100** | 0.409<br>0.372<br>0.415<br>0.417<br>0.440<br>**0.441** | **-7.94**<br>-5.86<br>-7.07<br>-7.5<br>-7.06<br>-7.08 | **0.667**<br>0.607<br>0.617<br>0.577<br>0.652<br>0.642 | X<br><br><br><br>X<br>X | X<br><br><br><br>X<br>X |
