## Supplementary Table S3 for "*In silico* study predicts a key role of RNA-binding domains 3 and 4 in nucleolin-miRNA interactions"

**Table S3:** NCL interacting miRNAs based on experimental evidence or predicted interaction with NCL.

| MicroRNA family/clusters | Interaction evidence/prediction | Cell/Tissue Type |
|---|---|---|
| **mir-15a, mir-16** | RNA immunoprecipitated with NCL-specific antibodies (RNA IP), decrease mature miRNA levels/increased Pri-miRNA levels upon NCL knockdown | HEK293 cells (human embryonic kidney) / MCF7 Breast cancer cells [15] |
| **mir-21, mir-221, mir-222, mir-103** | RNA IP, altered miRNA levels and miRNA target levels upon NCL depletion | HeLa cells / MCF7 Breast cancer cells [40] |
| **Clusters of mir-17-92, mir-15, mir-16, mir-221-222, 30, let-7, mir-155** | Predicted, transcriptomic analysis revealing altered miRNA levels upon NCL depletion | HeLa Cells [41] |
| **mir-93, mir-484** | Predicted, altered miRNA target levels upon NCL phosphorylation | Human Umbilical Vein Cells (HUVEC) [42] |
| **mir-223, mir-214, mir-146b, mir-199a-5p, mir-208b, mir-29a, mir-146a, mir-34a, mir-690, mir-582-5p, mir-135a, mir-145, mir-218** | Predicted, transcriptomic analysis revealing altered miRNA levels upon NCL overexpression | Transgenic mice [43] |
