## Supplementary Table S4 for "*In silico* study predicts a key role of RNA-binding domains 3 and 4 in nucleolin-miRNA interactions"

**Table S4: Evaluation and refinement of miRNA models.** All miRNA models were evaluated by MolProbity for structural integrity. For every category of evaluation, a lower score indicates a higher quality model.

| miRNA models | Clash score | Probable wrong sugar puckers | Bad backbone conformations | Bad bonds | Bad angles | Top ranked model |
|---|---|---|---|---|---|---|
| **miR-15a models** | | | | | | |
| RNACompstd | 22.49 | 1 | 15 | **0** | **0** | |
| RNAFold | 22.49 | **0** | 15 | **0** | **0** | |
| MCSYM | **17.24** | 1 | **4** | **0** | **0** | **X** |
| Centroid | 27.36 | 1 | 20 | **0** | **0** | |
| SPOT-RNA | 17.62 | 1 | 16 | **0** | **0** | |
| RNAStructure | 22.49 | 1 | 15 | **0** | **0** | |
| **miR-16 models** | | | | | | |
| RNACompstd | 20.05 | **2** | 17 | **0** | **0** | |
| RNAFold | 16.18 | **2** | 13 | **0** | **0** | |
| MCSYM | 13.01 | **2** | **5** | **0** | **0** | **X** |
| Centroid | 20.05 | **2** | 17 | **0** | **0** | |
| SPOT-RNA | **12.66** | **2** | 9 | **0** | **0** | **X** |
| RNAStructure | 16.18 | **2** | 13 | **0** | **0** | |
| **miR-21 models** | | | | | | |
| RNACompstd | 19.98 | **0** | 11 | **0** | **0** | |
| RNAFold | 21.29 | **0** | **9** | **0** | **0** | **X** |
| MCSYM | 20.85 | **0** | 7 | **0** | **0** | **X** |
| Centroid | 19.98 | **0** | 11 | **0** | **0** | |
| SPOT-RNA | 49.96 | 1 | **9** | 10 | 27 | |
| RNAStructure | **16.51** | 2 | 12 | **0** | **0** | |
| **miR-103 models** | | | | | | |
| RNACompstd | 19.32 | 4 | 24 | **0** | **0** | |
| RNAFold | 15.69 | 4 | 17 | **0** | **0** | |
| MCSYM | **14.49** | **1** | **9** | **0** | **0** | **X** |
| Centroid | 18.91 | 3 | 18 | **0** | **0** | |
| SPOT-RNA | 22.94 | 3 | 16 | **0** | **0** | |
| RNAStructure | 16.9 | 4 | 19 | **0** | **0** | |
| **miR-221 models** | | | | | | |
| RNACompstd | 25.07 | **2** | 28 | 8 | 6 | |
| RNAFold | 25.93 | **2** | 25 | 8 | 6 | |
| MCSYM | 70.66 | 4 | **18** | 17 | 44 | |
| Centroid | 20.23 | 3 | 27 | **0** | **0** | |
| SPOT-RNA | **17.09** | 3 | 21 | **0** | **0** | **X** |
| RNAStructure | 24.79 | **2** | 28 | 8 | 6 | |
| **miR-222 models** | | | | | | |
| RNACompstd | 17.69 | 2 | **14** | **0** | 1 | |
| RNAFold | 17.69 | 2 | **14** | **0** | 1 | |
| MCSYM | 18.54 | 2 | 22 | **0** | **0** | |
| Centroid | **14.27** | **1** | 20 | **0** | **0** | **X** |
| SPOT-RNA | 17.69 | 3 | 19 | **0** | **0** | |
| RNAStructure | 17.69 | 2 | **14** | **0** | 1 | |

| **mir-155 models** | | | | | | |
|---|---|---|---|---|---|---|
| RNAFold/RNAStruc | 13.58 | 1 | **12** | **0** | 0 | |
| Centroidfold | 18.91 | 1 | **10** | **0** | 0 | |
| MCSYM | 19.4 | 2 | **14** | **0** | 0 | |
| SPOTRNA | 18.43 | 0 | **10** | **0** | 0 | **X** |
