## Supplementary Figure S1 for "*In silico* study predicts a key role of RNA-binding domains 3 and 4 in nucleolin-miRNA interactions"

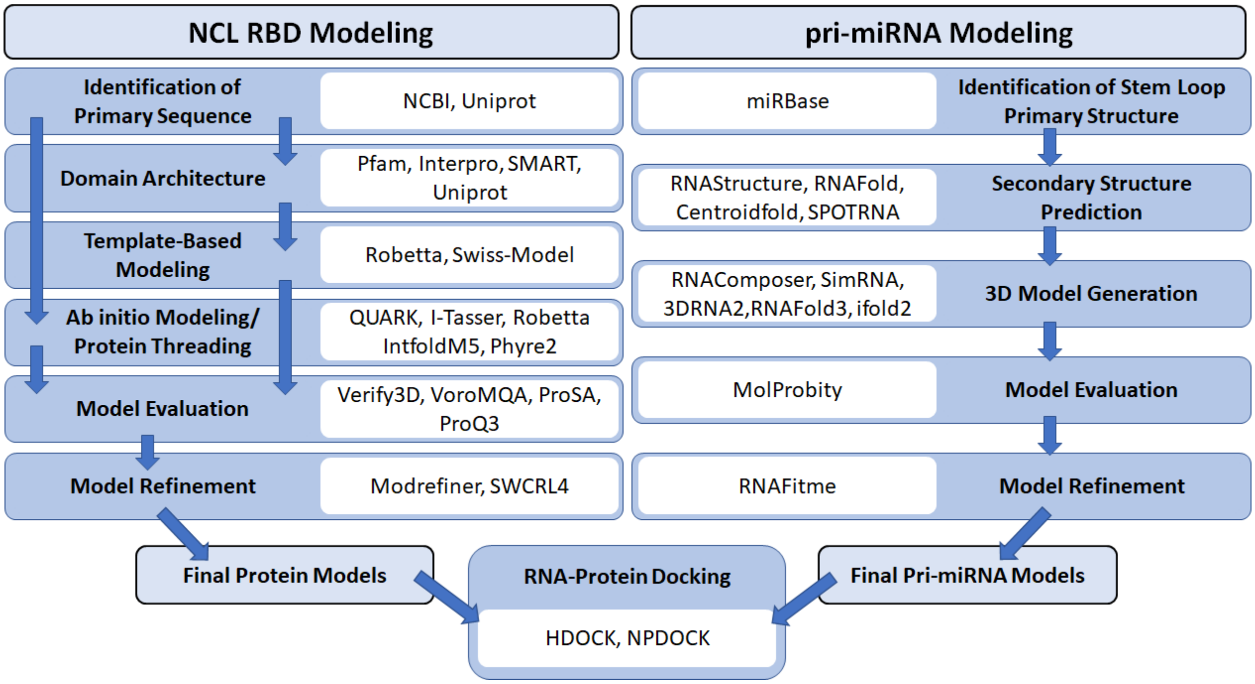


**Figure S1: The workflow and tools used in this study.** The schematic workflow devised to model the NCL-RBDs, miRNA and characterize their interactions. Blue boxes and arrows indicate sequential processing steps while white boxes denote the programs/tools used in the corresponding steps for protein and pri-miRNA modeling are indicated.
