## Supplementary Figure S2 for "*In silico* study predicts a key role of RNA-binding domains 3 and 4 in nucleolin-miRNA interactions"

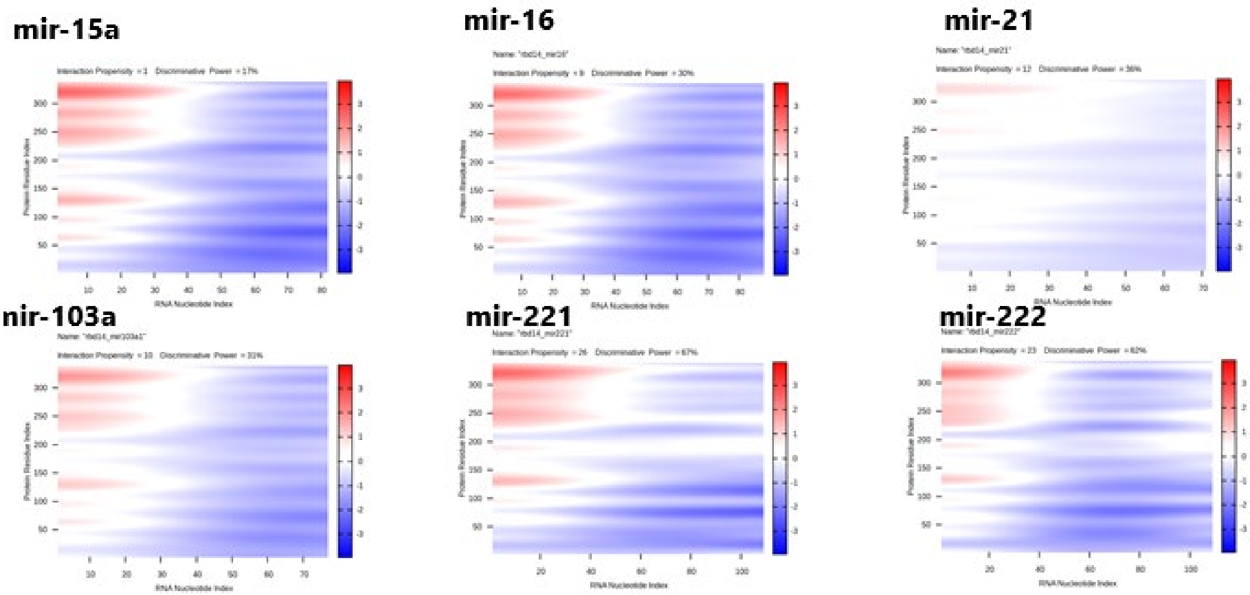


**Figure S2**. CatRAPID results of NCL RBD14 for all miRNA sequences used in this study. Red regions indicate a strong indicator for RNA binding propensity (the taller the red peaks, the greater the propensity). RBD1: 1-77 RBD2: 87-160 RBD3: 180-254 RBD4: 266-34
