## Supplementary Figure S3 for "*In silico* study predicts a key role of RNA-binding domains 3 and 4 in nucleolin-miRNA interactions"

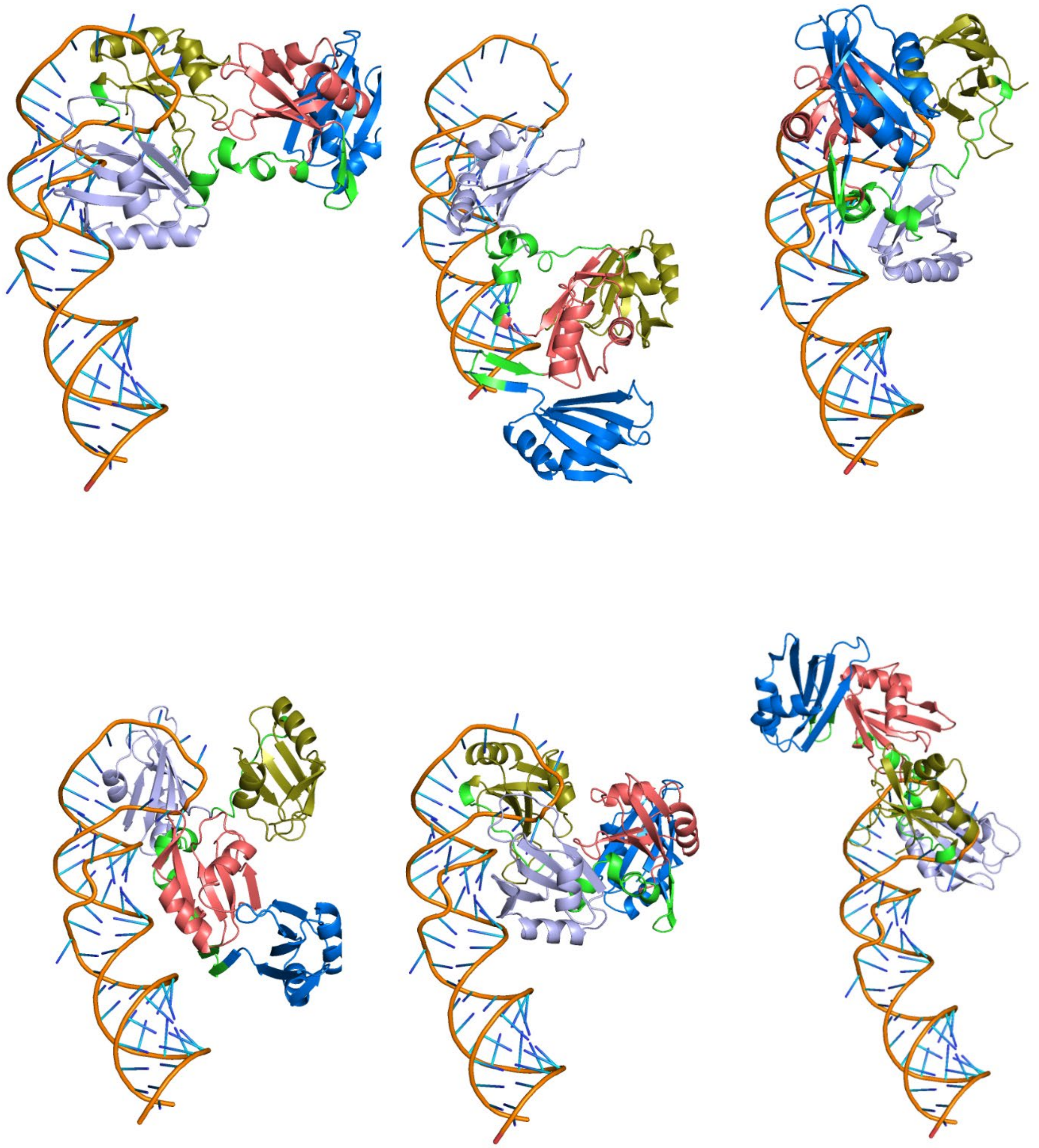

**Figure S3. Inconsistent interactions displayed by RBD14-mir155 docking results.** RBD1 (marine blue), RBD2 (deep salmon), RBD3 (light blue), RBD4 (deep olive), and miRNA (orange), linker region connecting RBDs (neon green)
