## Supplementary Figure S4 for "*In silico* study predicts a key role of RNA-binding domains 3 and 4 in nucleolin-miRNA interactions"

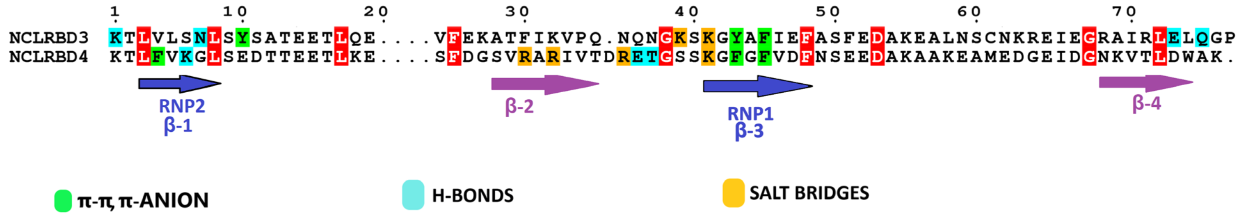


**Figure S4. MSA map of RBD34 predicted to be involved in NCL-miRNA interactions.** The type of interactions are color coded as indicated at the bottom of the figure.
