## Supplementary Figure S5 for "*In silico* study predicts a key role of RNA-binding domains 3 and 4 in nucleolin-miRNA interactions"

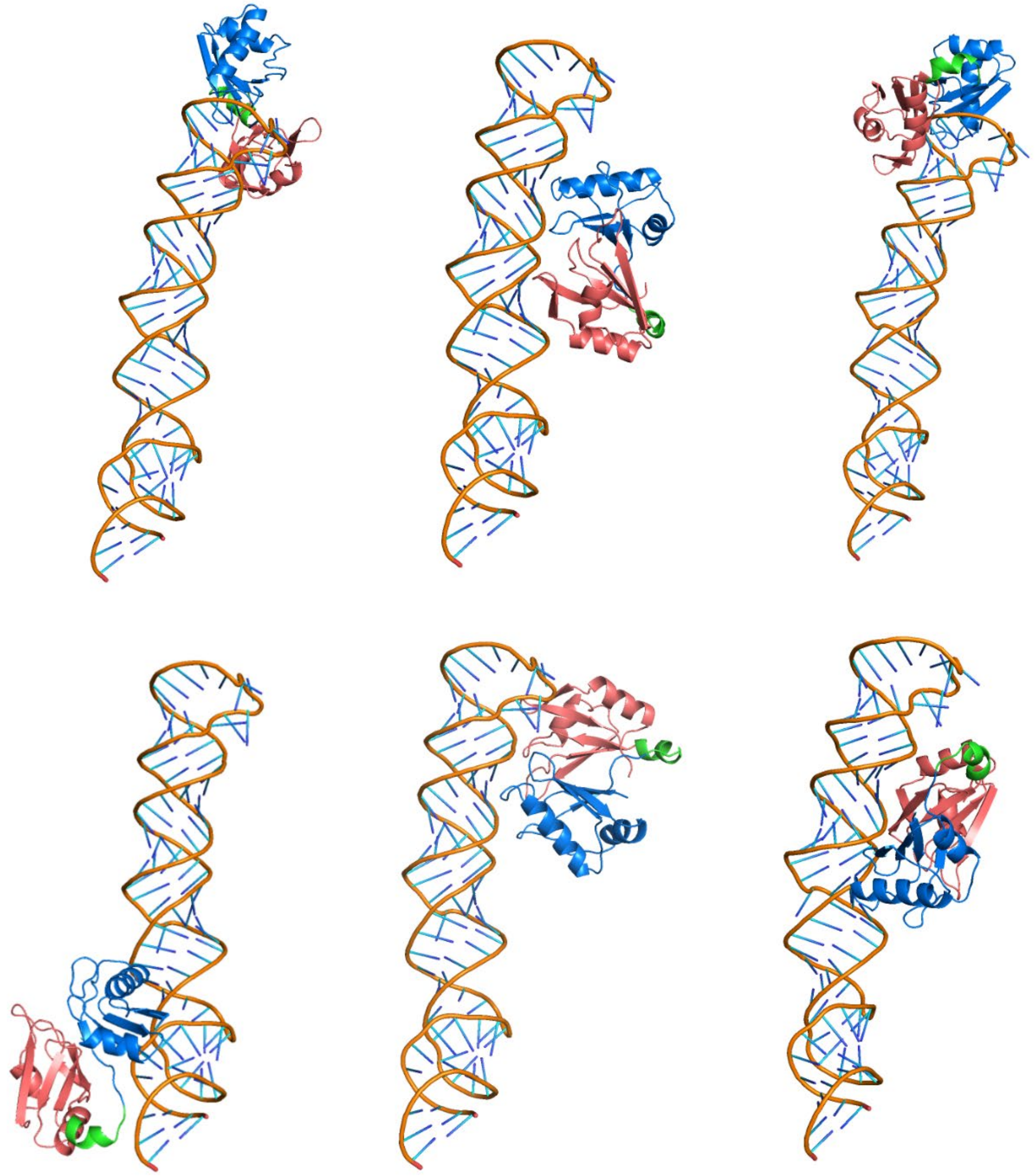

**Figure S5. Inconsistent scenarios of RBD1-2-mir16 interactions**. RBD1 (marine blue) , RBD2 (deep salmon), miRNA (orange), linker region connecting RBD12 (neon green)
